# Dancing the Nanopore limbo – Nanopore metagenomics from small DNA quantities for bacterial genome reconstruction

**DOI:** 10.1101/2023.02.16.527874

**Authors:** Sophie A. Simon, Katharina Schmidt, Lea Griesdorn, André R. Soares, Till L. V. Bornemann, Alexander J. Probst

## Abstract

**Background:** While genome-resolved metagenomics has revolutionized our understanding of microbial and genetic diversity in environmental samples, assemblies of short-reads often result in incomplete and/or highly fragmented metagenome-assembled genomes (MAGs), hampering in-depth genomics. Although Nanopore sequencing has increasingly been used in microbial metagenomics as long reads greatly improve the assembly quality of MAGs, the recommended DNA quantity usually exceeds the recoverable amount of DNA of environmental samples. Here, we evaluated lower-than-recommended DNA quantities for Nanopore library preparation by determining sequencing quality, community composition, assembly quality and recovery of MAGs.

**Results:** We generated 27 Nanopore metagenomes using the commercially available ZYMO mock community and varied the amount of input DNA from 1000 ng (the recommended minimum) down to 1 ng in eight steps. The quality of the generated reads remained stable across all input levels. The read mapping accuracy, which reflects how well the reads match a known reference genome, was consistently high across all libraries. The relative abundance of the species in the metagenomes was stable down to input levels of 50 ng. High-quality MAGs (> 95% completeness, ≤ 5% contamination) could be recovered from metagenomes down to 35 ng of input material. When combined with publicly available Illumina reads for the mock community, Nanopore reads from input quantities as low as 1 ng improved the quality of hybrid assemblies.

**Conclusion:** Our results show that the recommended DNA amount for Nanopore library preparation can be substantially reduced without any adverse effects to genome recovery and still bolster hybrid assemblies when combined with short-read data. We posit that the results presented herein will enable studies to improve genome recovery from low-biomass environments, enhancing microbiome understanding.

## Introduction

Metagenomics has expanded our knowledge of the diversity of Bacteria, Archaea, and Viruses across all environments on Earth [*c*.*f*., [1–3]]. Recovering MAGs has become a key practice to assess the taxonomic and functional diversity of microorganisms in various environments while circumventing the need to cultivate the respective microbes for genomic studies [4]. Genome-resolved metagenomics provides detailed insights into crucial microbially-driven roles across various ecological processes, such as nutrient cycling [5] or their interactions with other organisms [6]. The completeness and contamination of a reconstructed MAG are directly correlated with the accuracy of downstream genomic predictions, which can elucidate the metabolic potential [7, 8], evolutionary relationships [9], horizontal gene transfer [10], and other genomic traits [*c*.*f*., [11–13]]. Using short-read sequencing platforms, the first circularized, closed MAGs (cMAGs) were recovered about ten years ago [14, 15]. In September 2019, the number of publicly available cMAGs was 59 [16] compared to the thousands deposited MAGs in public databases [*c*.*f*., [17, 18]]. Using a hybrid approach of short Illumina reads and long Nanopore reads, Singleton *et al*. have recently been able to recover almost the same amount of circularized, complete MAGs (57) within one study and from the same ecosystem [19].

Oxford Nanopore Technologies (ONT) has emerged as powerful platform for metagenomics as long reads produced by this sequencing technology span large areas of genomes, covering otherwise problematic genomic regions. Examples for such regions are highly repetitive and/or conserved elements including multiple copies of the same transposable element in one genome [20], for which short read-based *de novo* assemblies are more likely to fail. Long-read metagenomics has successfully been used to for cMAG recovery from diverse samples, including activated sludge [21, 22], water bodies [23], sediment [24] and feces [25, 26]. However, all aforementioned ecosystems were of high biomass, providing sufficient DNA amounts for Nanopore sequencing.

ONT recommends using 1000 ng high-molecular weight (HMW) DNA as input for preparing sequencing libraries using the ligation sequencing kits SQK-LSK109 and SQK-LSK110 and loading between 5 and 50 fmol DNA library on a R9.4.1 Flow Cell. Since these DNA quantities and molarities are often difficult to extract from environmental samples, we investigated how Nanopore sequencing quality, stability of coverage distribution, assembly, and binning quality varies with reduced DNA input quantities. To this end, we sequenced a microbial standard of high molecular weight DNA reducing the amount of input DNA for library preparation from 1000 down to 1 ng in eight steps. Evaluating the results at multiple levels of a genome-resolved metagenomics pipeline (from reads to assemblies and MAGs) we demonstrate that the required amount for successful assembly and genome reconstruction can be significantly reduced. Based on these results, we recommend including Nanopore long reads in every assembly-based metagenomic study including those from low-biomass environments.

## Results

To determine the lower limit of DNA input for Nanopore sequencing, we generated 27 long read metagenomes (nine input levels, three replicates each), whereas the ZymoBIOMICS HMW DNA standard served as input material. This mock community was composed of seven bacterial strains and one yeast strain (https://files.zymoresearch.com/protocols/_d6322_zymobiomics_hmw_dna_standard.pdf [02.02.23]); the range of input varied from 1000 ng (recommended) down to 1 ng. Each input quantity was analyzed in triplicates and is herein reported with mass and replication number, *e*.*g*., 350_3 for 350 ng input material and replicate #3. The amount of DNA library loaded onto the Flow Cell was reduced from the recommended amount of 4.7 to 9.88 fmol prepared from 1000 ng down to 0.136 fmol prepared from 10 ng DNA input; five sequencing libraries were below the detection limit of the Qubit DNA HS assay but still successfully sequenced (Table S1). We aimed for 1 Gbp of raw sequencing depth per metagenome; down to 50 ng input material, 1 Gb was achieved for all but three metagenomes (350_2, 200_3, 100_3). For the nine metagenomes with less DNA input, only one sequencing run with 1 ng input material (1_1) met the target output (sequencing depth is before and after quality control is provided in Table S2). We assessed the results of the different input levels based on read quality, stability of the coverage distribution across the individual microbial genomes, the assembly quality, and the quality of *de novo* reconstructed bacterial genomes. In addition, we leveraged publicly available Illumina data of the mock community to determine if long-reads from low input material can bolster hybrid assemblies of microbial communities [27].

### Sequencing quality remains stable irrespective of DNA input quantity

The Q-Score of the generated reads varied around 12.61 (± 1.11) and remained stable across the entire dilution series (Figure 1A). No difference in read quality was evident between the two sequencing kits used herein (SQK-LSK109/110), rendering all metagenomes comparable with each other. The length of nanopore reads, represented using read length N50, had a mean of 19,106 bp (± 4991 bp) in the range of 1,000 ng down to 50 ng (Figure 1B). For lower input levels, we observed a significant decrease (U-test, p-value <0.001) of the read length N50 down to an average of 3883 bp (± 957 bp) (Figure 1B) compared to the 1000 ng – 50 ng range. The read mapping accuracy by read-to-reference alignments varied between 94.68% and 99.98% for all samples, demonstrating fairly consistent sequencing quality irrespective of input DNA quantity with the exception of an improved mapping rate of 1 ng samples compared to 1000 ng (Kruskal-Wallis p-value 0.0614 across all samples, post-hoc Dunn’s test p-value 0.0298 for this comparison only; Figure 1C).

**Figure 1:**
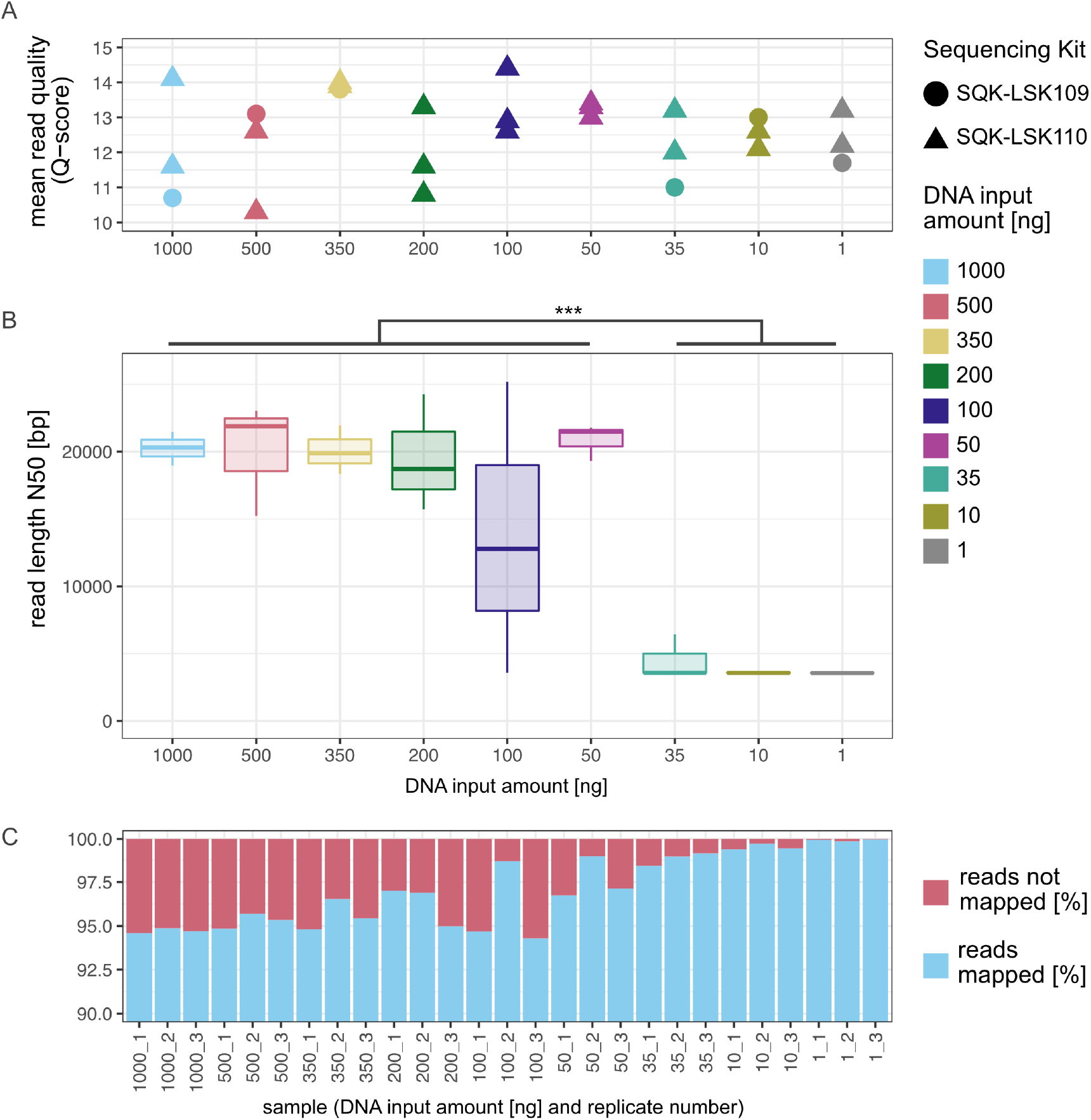
Sequencing summary. A) Mean read quality (Q-Score) of all Nanopore metagenomes differentiated by the used sequencing kit. B) Boxplot showing how the read length, represented by the N50, behaves over the course of the input reduction. The read length N50 decreases significantly when sequencing less than 50 ng. To test for significance the values for the N50 were grouped into ≥ 50 ng and ≤ 35 ng. Both groups are not normally distributed (Shapiro Wilk test; p-value ≥ 50 ng 4.136e-07; p-value ≤ 35 ng 0.004872), with a subsequent Wilcoxon Rank test the significance was verified (W = 1, p-value = 4.32e-05). C) Proportions of sequencing reads that map to reference genomes changes. The mapping rate varied between 94.29 and 99.98 %.

### Community composition remains stable down to 50 ng DNA input

To compare the coverage or percent relative abundance of the eight species in the metagenomes with their theoretical abundance in the community standard, we mapped all reads to the reference genomes using minimap2 [28]. We were able to detect all eight species included in the community standard in each of the 27 metagenomes (Figure 2A). Except for one outlier (100_3), all samples down to an input of 50 ng showed a significant Pearson correlation (p-value < 0.05) between the determined percent relative abundances of the species and the theoretical abundance of the individual species (correlation and p-values in table S3). When further reducing the DNA input, the relative abundance of *Salmonella enterica* and *Escherichia coli* increased tremendously to more than 95 % in combination. A principal coordinate analysis, PCoA, of the relative abundance distribution between samples and theoretical composition shows clustering of replicates for inputs ranging from 50 to 1000 ng, with the distance between replicates increasing inversely with input amount (Figure 2B). Metagenomes generated from 1 ng - 35 ng input material were strongly separated along the first axis of the PCoA (one 100 ng library represents an outlier, see above), which agrees with the strong distortion of the community below 100 ng input material observed in the bar plot (Figure 2A). Surprisingly, in the range of 1 to 35 ng of input DNA, distances between replicates decrease significantly, suggesting high reproducibility in this input range.

**Figure 2:**
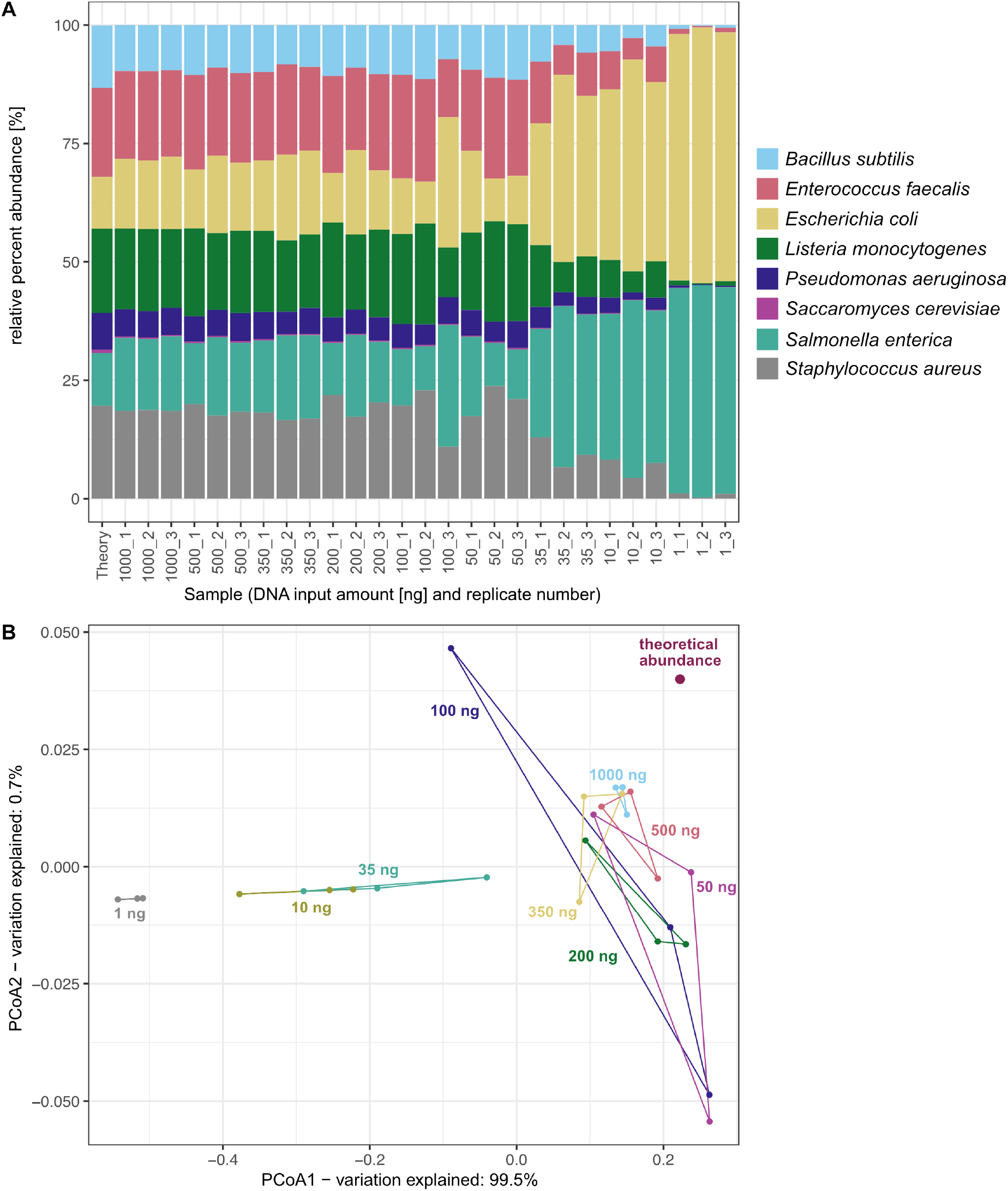
Abundance analysis. A) Relative abundance and taxonomic assignment based on mapping Nanopore sequences to each of the eight reference genomes. B) PCoA based on Bray-Curtis dissimilarities of percent relative abundance of species calculated from read coverage in comparison to the theoretical composition of the sequenced microbial standard (ZYMO).

### A read of length is a joy forever: Nanopore reads derived from down to 1 ng DNA input improve hybrid assemblies

For Nanopore reads assembled using metaFlye, the average N50 of the assembly was 3.89 Mb (± 1.46 Mb) down to an input level of 50 ng. Fragmentation of the assemblies was the greatest for libraries generated from low DNA quantity (1 ng - 35 ng), while high-input libraries showed the greatest N50. However, Nanopore reads of replicates 2 and 3 corresponding to 1 ng input could hardly be assembled, *e*.*g*., the assembly of replicate 2 of 1 ng only had a total size of 31,902 bp distributed over 4 contigs (Table S4). To determine if low input material could still improve hybrid assemblies, we assembled our Nanopore sequencing data with publicly available Illumina data [27] of the mock community for all 27 metagenomes using hybridSPAdes [29]. The N50 of the hybrid assemblies was 1.06 Mb (± 0.259 Mb) for input levels down to 50 ng. At lower amounts, the N50 dropped drastically compared to assemblies of higher input material to an average of 0.375 Mb (± 0.246 Mb). The lowest N50 of a hybrid assembly was 136 kb, achieved with Nanopore reads obtained from 1 ng DNA, but still higher than that of a short-read only assembly (120 kb; Figure 3A).

**Figure 3:**
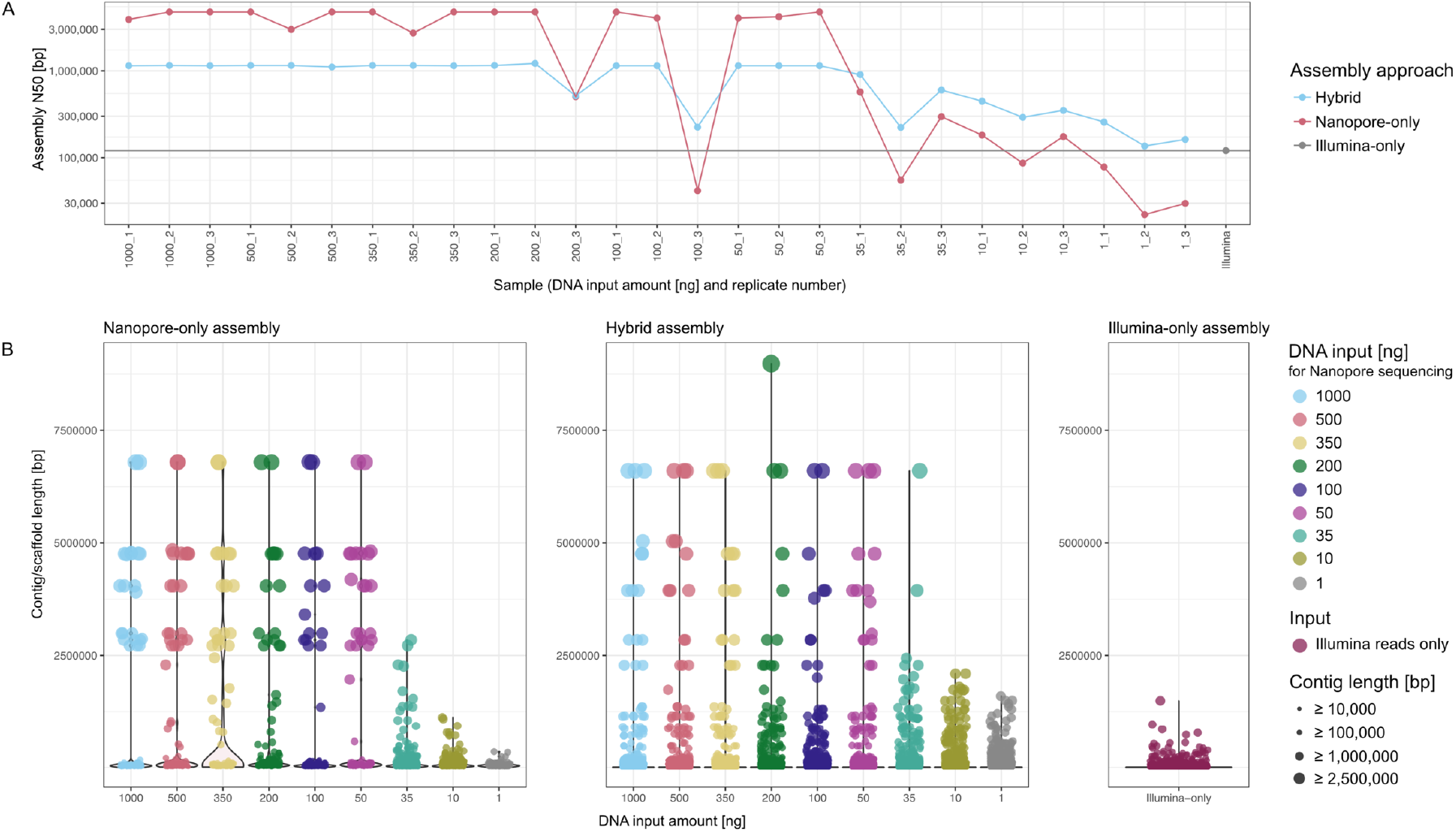
Comparison of undertaken assembly approaches. A) Representation of the N50 across Nanopore-only, hybrid and Illumina-only assemblies. B) Overview showing the contig/scaffold to length distribution faceted by assembly approach.

While the GC content of the hybrid and Illumina-only assembly was very constant (on average 47.41% (± 0.04 %), the more variable GC content of Nanopore assemblies was 46.87% (± 3.11 %). Down to 50 ng input, the GC content of the Nanopore assemblies is at 48.38% (± 1.45%), and then decreased to 43.84 % (± 3.40%). Nanopore reads showed less variation in GC content than the assemblies (44.05 ± 1.01 %; Figure S1).

Analyses of contig/scaffold lengths support the conclusions of N50 analyses. Using only Nanopore reads down to 50 ng input material, contig lengths that corresponded to the respective expected genome sizes of mock community species assembled. For example, in twelve of the 27 assemblies, the greatest contig was 6.79 Mb in length, which is the genome size of *Pseudomonas aeruginosa*. With hybrid assemblies, scaffolds of around 6.6 Mb could be detected in 17 of 27 assemblies. This approach also enabled the reconstruction of long scaffolds at 35 ng input material, which was not possible for short or long reads alone (3.9 and 6.6 Mbps respectively). Indeed, the longest scaffold of the short-read only assembly has a length of 1.49 Mb, which is substantially shorter than the smallest genome in the mock community (*Staphylococcus aureus*, 2.730 Mbps). The longest scaffolds resulting from hybrid assemblies fed with Nanopore reads from 1 ng and 10 ng input DNA ranged between 1.50 Mb and 2.09 Mb (highest for 1 ng 1.59 Mbps), which is still greater than short-read assembly alone (Figure 3B).

### Recovery of near-complete MAGs down to 35 ng input material

After assembly, the Nanopore-only metagenomes were manually binned using uBin [30]. uBin uses 51 universal bacterial single-copy genes as markers [31]. As depicted in Figure 4A, recovery of near complete genomes (**≥** 95% completeness, **≤** 5% contamination) was successful down to 50 ng of input material. Remarkably, we could also bin 4 high quality MAGs with more than 95 % completeness from replicate 35_3, *i*.*e*., with 35 ng input material only. Analysis of Average Nucleotide Identity (ANI) and length consensus—a metric comparing how well the sequences of a MAG and reference sequence align to each other—confirmed the high quality of the recovered MAGs (Figure 4B). These comparisons showed that even down to 10 ng input, bins with a length consensus > 75% and an ANI of > 99.7% could be obtained for Nanopore-only metagenomes (Figure 4B).

**Figure 4:**
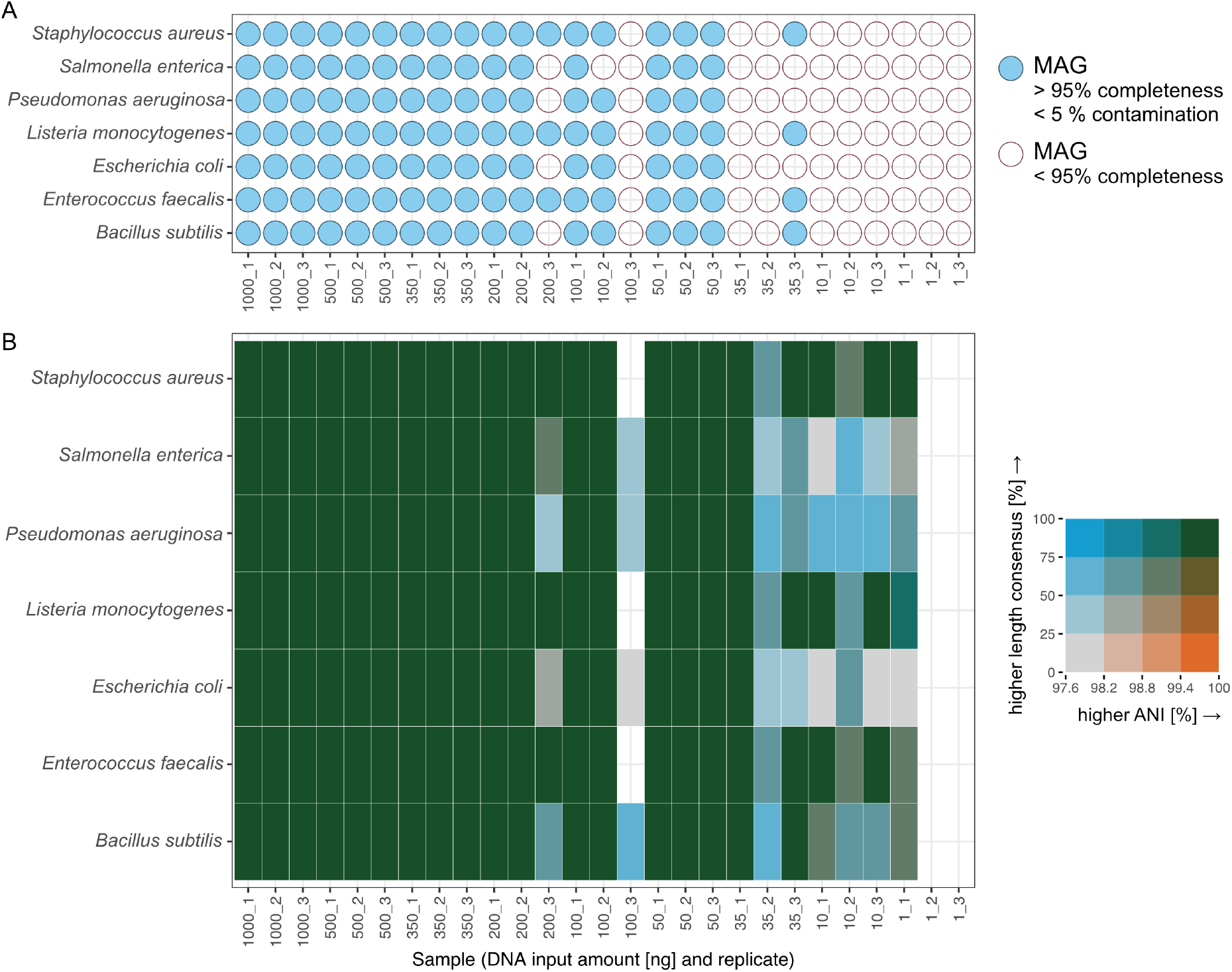
Quality of metagenome assembled genomes. A) Bubble plot showing which genomes were near-full length recovered based on single copy genes. B) Bivariate heatmap showing the relationship between length consensus and ANI per bin. The increments of the legend were distributed equally on both axes.

## Discussion

In this study, we report how lower than recommended DNA input affects Nanopore long-read metagenomics by comparing different levels of metagenomic analysis from read QC to retrieval of MAGs. Many previously published Nanopore-based metagenomics studies lack information about the quantity of DNA used for library preparation [19, 24–26, 32]. To the best of our knowledge, other published metagenomic studies quantified the amount of input DNA and met or exceeded the recommended amount of 1000 ng DNA for input material. These studies covered various high-biomass environments including human oral cavity [33, 34], enrichments of artificially contaminated irrigation water [35], groundwater aquifer [36], sand filters of a drinking water treatment plant [37], and activated sludge [21] with up to 4.5 µg of DNA for library preparation. Noteworthy, Chen *et. al*. spiked environmental DNA with DNA of a known bacterium as the authors described their DNA amount as insufficient for Nanopore sequencing [23]. In similar fashion, the so-called CarrierSeq protocol includes the addition of 1000 ng of Lambda phage DNA to the target DNA, which could be metagenomic DNA for sequencing; in such an approach Mojarro et al. used 0.2 ng *Bacillus subtilis* DNA as the target, which resulted in only 777 Nanopore-reads (out of 718,432) that mapped to the target genome of *B. subtilis* [38]. In a study of low-biomass Mars analog soil, CarrierSeq was used for astrobiological investigations also resulting in limited amount of sequence information [39]. With the exception of the aforementioned study, all studies listed above utilized the recommended quantity of DNA from microbiomes that are of high biomass or easily accessibility for repeated sampling. However, there are numerous examples of low-biomass ecosystems, for which DNA extractions do not yield the recommended DNA input for ONT library preparation and excessive sampling is difficult.

While the most prominent ecosystems on Earth (soil, ocean) usually harbor high biomass, smaller ecosystems with low biomass are far more diverse and numerous and yet important for ecosystem services and human health. The air microbiome, for instance, is an ecosystem that constantly and globally surrounds humans, influences the Earth’s surface but remains little explored regarding the genetic diversity resolved at species level [40]. The gaseous state, and the low biomass, atmospheric turbulences, temperature differences and day and night shifts complicate sampling [40, 41]. Consequently, we suggest that low-input Nanopore sequencing combined with short reads in a hybrid assembly as presented herein could substantially enhance the study of the air microbiome. In similar fashion, research on glacier and desert microbiomes could be facilitated via hybrid assemblies.

Glacial ice for example, contains minute microbial cell concentrations when compared to many other environments, making low-biomass metagenomics of their unique microbial communities necessary [42, 43]. This is also the case for desert soils, where little DNA can be recovered, and resulting genomes from short-read assemblies are generally highly fragmented [44]. Contamination control of spacecraft assembled in cleanrooms is of uttermost importance to protect the integrity of life-detection missions destined to other celestial bodies [45]; however, cleanroom environments are of low-biomass, and application of metagenomics to cleanroom samples usually requires whole genome amplification prior to sequencing [46] distorting community compositions [47] and thus hampering the recovery of genomes from metagenomes. In addition, research on environments for which the matrix strongly hinders nucleic acid extraction could also benefit from approaches developed for low-biomass environments. For instance, chemical processes (*e*.*g*., induced by pH) can lead to DNA hydrolysis, denaturation or depurination [48], or nucleic acids are captured by adsorption effects [49]. Furthermore, there are ecosystems where quantity as well as access is restricted (*e*.*g*., the surface of the human eye [50]); at the same time, high spatial resolution of high-biomass environments like soil could also be achieved via lowering input material. In sum, there are numerous examples of ecosystems on Earth that could substantially benefit from long-read low-input metagenomics to bolster our understanding of microbial diversity and evolution.

Based on our results, we recommend adding Nanopore sequencing for metagenomic studies at each DNA concentration investigated, *i*.*e*., from 1000 ng down to 1 ng. The recovery of circularized, closed genomes and the improvement in scaffold length clearly shows that hybrid approaches of short and long reads should be the current standard. To close genomes of, *e*.*g*., slow growing prokaryotic isolates, co-cultures or low complex enrichment cultures using Nanopore sequencing 35-50 ng DNA are sufficient as starting material.

## Outlook

Although our study has successfully demonstrated that low DNA input Nanopore long-read metagenomics is possible, we acknowledge certain limitations in terms of extrapolating our findings to very complex environmental samples or new ONT chemistries that have become available while the study was executed or will become available in the future. For instance, with the new Q20+ chemistry, the recommended input amount for a library preparation is still 1000 ng of HMW DNA or > 100 ng of fragmented DNA. Recently, the recommended amount of final DNA to be loaded onto the Flow Cells has remained constant or even decreased compared to the previously used combination of LSK109/110 and R9 Flow Cells, which is promising for the future application of Nanopore sequencing for exploring low-biomass ecosystems. Previously, 5-50 fmol DNA library were recommended, but at the moment of writing this paper (February 2023), Oxford Nanopore Technologies recommends only 5-10 or 10-20 fmol for the new chemistries. Another interesting development that was recently announced by ONT for handling small DNA samples is Library Recovery. The DNA library is sequenced on one Flow Cell and as soon as its sequencing performance decreases or ends, the library is aspirated out of the original Flow Cell with a pipette tip and transferred to a new, freshly primed Flow Cell to continue sequencing (https://community.nanoporetech.com/docs/prepare/library_prep_protocols/library-recovery-from-flow-cells/v/lir_9178_v1_revb_11jan2023 [18.01.23]). Exploring these new techniques in combination with low DNA input for library preparation will be a challenging but also promising task with the goal to further explore biodiversity and genetic content of low-biomass environments.

## Materials and Methods

### Microbial Community Standard

The ZymoBIOMICS HMW DNA Standard #D6322 (Zymo Research, USA) was used to determine the input limits for Nanopore gDNA sequencing without the need for whole-genome amplification. The DNA standard is composed of eight microbial species: seven bacteria and one yeast (Table S5). Used reference genome sequences are provided by Zymo and available at https://s3.amazonaws.com/zymo-files/BioPool/D6322.refseq.zip [07.12.2022].

### Library preparation and QC

The input DNA amount of the standard was reduced stepwise from the recommended 1000 ng to 1 ng and included the following quantities of DNA: 1000 ng – 500 ng – 350 ng – 200 ng – 100 ng – 50 ng – 35 ng – 10 ng – 1 ng. DNA amounts in the dilution series were verified via Qubit. Each input quantity was analyzed in triplicates reported with mass and replicated number, *e*.*g*., 350_3 for 350 ng input material and replicate #3.

Library preparation was performed using the SQK-LSK109 and SQK-LSK110 sequencing kits (Oxford Nanopore Technologies, UK) with minor deviations from the manufacturer’s instructions: Both clean-up steps with AmPure XP beads (Beckman Coulter, USA) were extended by 5 min (10 min in total). To enrich for long fragments, AmPure XP beads were washed using the long fragment buffer (LFB) in the respective step. Additionally, elution of the DNA library from the AmPure XP beads was performed at 37°C as recommended for HMW DNA. DNA concentration and quality of prepared libraries were determined using Qubit 3.0 fluorometer (Thermo Fisher Scientific, USA) with the dsDNA HS array and Agilent TapeStation genomic DNA screen tapes. The maximum possible amount (12 µL) of DNA library was always loaded onto the Flow Cells. Prepared libraries were stored at -80°C if not subsequently sequenced.

### Nanopore Sequencing

Sequencing was performed using a MinION(tm) Mk1B (ONT) equipped with FLO-MIN106D Flow Cells. Sequencing runs were supervised by MinKNOW v21.02.1. Sequencing runs were stopped manually after achieving 1 Gb per sample or if even after > 12 hours the sequencing output had reduced so much that we did not expect to reach 1 Gb.

### Basecalling and read QC

Generated Nanopore raw reads were basecalled using Guppy v 6.1.3 in its super-accurate mode enabled using the dna_r9.4.1_450bps_sup.cfg model (https://community.nanoporetech.com/downloads [21.12.22]). Guppy basecalling in super-accurate mode automatically excluded reads with a Q-Score lower than 10. Statistics about the sequencing run and the sequencing reads were acquired using Nanoplot v 1.39.0 using the sequencing summary file generated by Guppy as input [51]. Basecalled Nanopore reads were filtered with Filtlong v 0.2.0 (https://github.com/rrwick/Filtlong [11.08.22]) to remove reads shorter than 2000 bps using –min_length 2000.

### Estimation of sequencing coverage

In order to estimate the coverage and the relative abundance of the individual species, respectively, Nanopore reads were mapped to the provided reference genome sequences using minimap2 v. 2.24-r1122 using the option --ax map-ont, --secondary=no and --sam-hit-only, whereas to quantify unmapped reads the --sam-hit-only flag was omitted [28]. Samtools v. 1.10 was used to manipulate resulting SAM files [52].

### Long read metagenomic assembly and binning

Long reads were assembled using Flye v. 2.9-b1768 with --meta and --nano-raw options enabled [53]. Open reading frames were predicted using prodigal v.2.6.3 [54] in its meta mode and annotated using DIAMOND v.0.9.9 [55] by searching against UniRef100 (e-value cutoff: 0.00001) [56]. Nanopore reads were mapped against the contigs to determine contig coverage using minimap2 as described above alongside uBin helper scripts to determine length and GC content per contig for manual binning (https://github.com/ProbstLab/uBin-helperscripts [18.01.23]). Manual binning and estimation of completeness and contamination of the genome bins were carried out using uBin [30]. Plasmids were not included in the bins.

### Assessment of MAG quality

The average nucleotide identity (ANI) between reference genomes and bins was calculated using fastani v.1.33 [57]. Agreement of reference genome and reconstructed MAG was determined by aligning them to each other using compare-sets (https://github.com/CK7/compare-sets [18.01.23]). For Assessing ANI and length consensus only the chromosomal genome was considered, *i*.*e*., plasmids were excluded from the reference genomes for this step.

### Illumina assembly and hybrid assembly

Illumina reads were taken from Sereika *et. al*. deposited at ENA with the BioProject ID RJEB48692 [27]. Quality control of Illumina reads was performed using BBduk (Bushnell B. – sourceforge.net/projects/bbmap/ [21.12.22]) and Sickle [58]. Reads were assembled using metaSPAdes 3.15.4 [59]. Hybrid assemblies, *i*.*e*. combinations of Illumina reads and Nanopore reads, were performed using hybridSPAdes3.15.4 with the --nanopore option [29]. Statistics like N50 and GC-content of all assemblies were assessed using SeqKit v.2.3.0 [60].

### Data visualization

Statistical evaluation and data visualization was done in R [61] using the packages tidyverse [62], ggplot2 [63], ggalt [64], ggnewscale [65], rcartocolor [66], ape [67] and biscale [68].

## Data availability

All raw sequencing data generated in this study will be deposited and made available at SRA upon publication.

## Acknowledgements

We thank Sabrina Eisfeld and Ines Pothmann for laboratory maintenance and Ken Dreger for server administration and maintenance. Maximiliane Ackers is acknowledged for administrative support.

## Financial Support

This study was funded by the German Federal Ministry of Education and Research within the project “MultiKulti” (BMBF funding code: 161L0285E).

## Author Contributions

Sophie A. Simon: Conceptualization, Methodology, Formal Analysis, Investigation, Project administration, Visualization, Writing – Original Draft

Katharina Schmidt: Investigation, Formal Analysis

Lea Griesdorn: Investigation

André Rodrigues Soares: Visualization, Writing – Review & Editing

Till L. V. Bornemann: Provided software, Writing – Review & Editing

Alexander J. Probst: Conceptualization, Methodology, Supervision, Project administration, Writing – Original Draft, Funding acquisition

## Conflict of interest

The authors declare that there are no conflicts of interest.

## Notes

### Competing Interest Statement

The authors have declared no competing interest.

