## Supplements for "Dancing the Nanopore limbo – Nanopore metagenomics from small DNA quantities for bacterial genome reconstruction"

List of contents:

1. Supplementary Tables
2. Supplementary Figure

### 1. Supplementary Tables

All tables are provided as a separate file <supplement_tables.xlsx>

**Table S1** | Information about Library Preparation and sequencing runs. The given concentration of the DNA library can be higher than the stated input DNA amount as the control DNA “DNA CS” was added during library preparation as recommended.

**Table S2** | Read statistics including total sequenced bases, length, and quality before and after quality control. We aimed for 1 Gb of raw sequencing data. Sequencing runs were manually stopped after 1 Gb was reached or sequenced until a plateau was reached.

**Table S3** | Pearson correlation between the theoretical percent relative abundance given in the information about the used ZYMO HMW DNA Standard and the measured percent relative abundance obtained.

**Table S4** | Assembly statistics for Nanopore-only, hybrid and illumina-only assembly.

**Table S5** | Composition and description of the ZymoBIOMICS HMW DNA Standard. Information taken from the instruction manual v 1.0.0 provided by Zymo.

### 2. Supplementary Figure

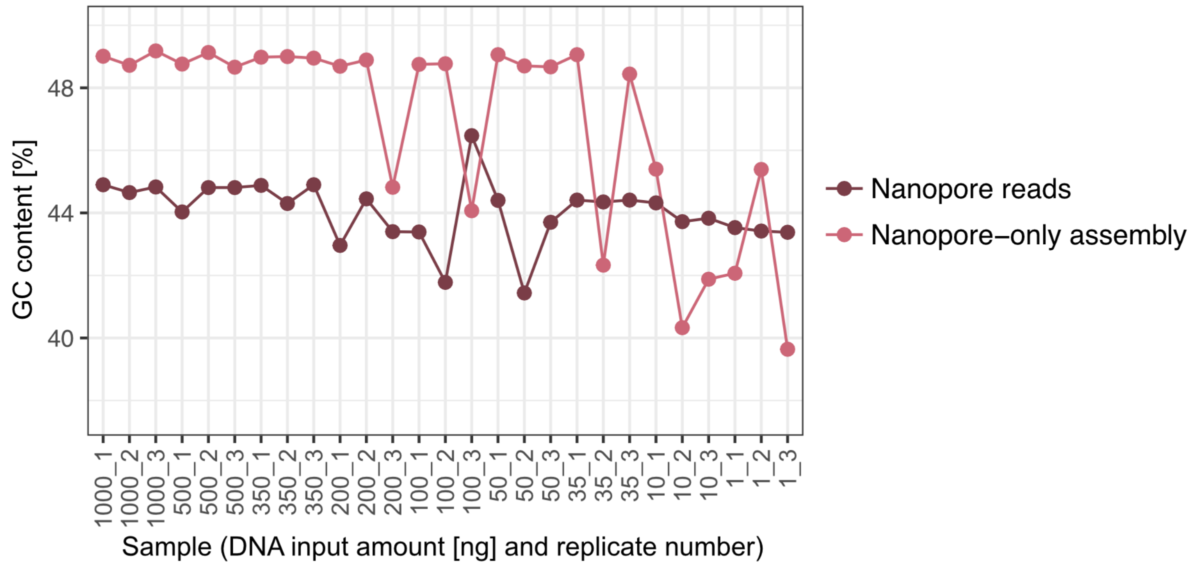


Figure S1 | GC-content [%] assessed for Nanopore Reads and Nanopore-only assemblies per sample. The GC-content varies distinctly for assemblies of less than 50 ng DNA input, while the GC content of the reads itself remains fairly stable. The GC content of the yeast *Saccharomyces cerevisiae* is given as 38.3% (https://files.zymoresearch.com/protocols/_d6322_zymobiomics_hmw_dna_standard.pdf, [26.01.23]), since it is not assembled by metaFlye, it is expected that GC content of the reads lower than of the assemblies.
